# Membrane contact site resident PTP1B limits superoxide production by suppressing a Syk-Shc1-Phagocyte Oxidase relay

**DOI:** 10.64898/2026.03.17.711902

**Authors:** Minhyoung Lee, Haggag S. Zein, Mahlegha Ghavami, Kuiru Wei, Murtaza Lokhandwala, Kaitlin Chan, Leanne Wybenga-Groot, Michael F. Moran, Gregory D. Fairn

**Affiliations:** Department of Biochemistry, University of Toronto, Toronto, Ontario, Canada; Department of Pathology, Dalhousie University, Halifax, Nova Scotia, Canada; Department of Biochemistry and Molecular Biology, Dalhousie University, Halifax, Nova Scotia, Canada; Department of Molecular Genetics, University of Toronto, Toronto, Ontario, Canada; Program in Cell & Systems Biology, and Program in Molecular Medicine, Hospital for Sick Children, Toronto, Ontario, Canada; SPARC BioCentre, Hospital for Sick Children, Toronto, Ontario, Canada

**Keywords:** Membrane contact site, NOX2, superoxide, phagocytosis, PTP1B, Syk kinase, Shc1

## Abstract

Phagocytosis is a specialized endocytic process used by macrophages and dendritic cells to engulf particles, which requires coordinated signaling cascades, cytoskeletal remodeling, and assembly of antimicrobial machinery to eliminate pathogens. During Fc γ receptor (FcγR)-mediated phagocytosis, dynamic actin depolymerization at the base of the phagocytic cup creates permissive conditions for endoplasmic reticulum-plasma membrane (ER-PM) membrane contact sites (MCS) to form. We demonstrate that the ER-resident protein tyrosine phosphatase PTP1B localizes to newly formed or expanded ER-PM MCS during phagocytosis and dephosphorylates Syk. Using TIRF microscopy with MCS residents, including MAPPER, STIM1, and E-Syts, we show that actin clearance allows ER proteins to approach the plasma membrane. PTP1B colocalizes with FcγRs in actin-cleared zones and physically interacts with Syk, a critical mediator of phagocytic signaling. Loss of PTP1B led to sustained Syk hyperphosphorylation without affecting phagocytosis. However, the PTP1B-deficient cells showed a ≍3-fold increase in NADPH oxidase 2 (NOX2)-mediated superoxide production. Using unbiased proteomics, we identified the adapter protein Shc1 as a critical intermediate linking Syk phosphorylation to NOX2 activation. Shc1 phosphorylation during phagocytosis is dependent on Src family kinases and Syk, while genetic ablation of *SHC1* reduced superoxide production by ≍40%. Proximity ligation assays reveal enhanced Shc1-p47phox interactions in PTP1B-deficient cells during phagocytosis. These findings establish an SFK-Syk-Shc1-NOX2 signaling axis that PTP1B negatively regulates at MCS between the ER and the forming phagosome, providing new mechanistic insights into antimicrobial responses during phagocytosis.

## Introduction

Phagocytosis is a fundamental cellular process by which professional phagocytes, such as macrophages and neutrophils, recognize, engulf, and destroy microbes and cellular debris, including apoptotic bodies^1–3^. Phagocytosis is a complex receptor-mediated endocytic process that requires exquisite coordination of multiple signaling pathways, cytoskeletal rearrangements, extensive membrane remodeling, and the assembly of microbicidal machinery on nascent phagosomes^4, 5^. Dynamic changes in the actin cytoskeleton are critical to support the internalization of phagocytic prey. First, actin polymerization at the tips of advancing pseudopods is essential to engulf particles. Second, coordinated removal of filamentous actin at the base of the phagocytic cup enables robust focal exocytosis to supply the membrane required to form the nascent phagosome^6, 7^. In addition to the uptake of target particles, microbial threats must also be destroyed. The phagocyte NADPH oxidase (NOX2) is central to antimicrobial defense by producing superoxide anion, which is converted to hydrogen peroxide^8, 9^. NOX2 is a multi-subunit enzyme composed of membrane-embedded components, gp91 and p22phox, and the recruitment of cytosolic components, p40phox, p47phox, and p67phox together with the small G-proteins Rac1 or Rac2^10, 11^. Together, the engulfment of prey and the antimicrobial actions in the phagosome effectively eliminate a wide range of microbes.

Fc γ receptor (FcγR)-mediated phagocytosis remains the most widely studied and best understood phagocytosis model^12^. Much of the signal transduction associated with FcγR-mediated phagocytosis is driven by phosphotyrosine-based signaling, with secondary signals produced by phosphatidylinositol 4,5-bisphosphate (PI4,5P_2_) hydrolysis, resulting in the generation of second messengers inositol trisphosphate (IP_3_) and diacylglycerol (DAG). These second messengers increase cytosolic calcium and activate protein kinase C isoforms^13^. Mechanistically, FcγR-mediated phagocytosis begins when IgG antibody-opsonized particles engage surface FcγRs, triggering activation of Src family kinases (SFKs) that phosphorylate immunoreceptor tyrosine-based activation motifs (ITAMs) within the FcγR cytoplasmic tails ^14, 15^. This creates docking sites for the Src Homology 2 (SH2) domain containing spleen tyrosine kinase (Syk), whose recruitment and activation are critical for downstream signaling events, including actin polymerization, membrane extension, and NOX2 assembly^4, 6, 16^.

ER-PM Membrane Contact Sites (MCS) are regions within cells where the ER is within 10-30 nm of the PM, and are critical hubs for cellular signaling and lipid homeostasis^17, 18^. These ER-PM MCS are stabilized by tethering proteins that bridge the two membranes and serve as platforms for calcium signaling, non-vesicular lipid transfer, and the localized regulation of PM lipid composition. Recent work has revealed that ER-PM contacts are dynamic structures that can be remodeled in response to cellular stimulation^19, 20^, suggesting they may play active roles in signal transduction beyond their established functions in lipid metabolism.

The intersection of ER-PM MCS biology with phagocytic signaling represents a minimally explored frontier in understanding innate immunity. Proteomics-based studies examining isolated phagosomes identified numerous ER proteins^21, 22^, which likely reflect ER-phagosome MCS rather than ER fusion with forming or maturing phagosomes^7, 23^. Indeed, evidence suggests that the non-fusogenic ER-resident SNARE protein Sec22b and syntaxin1 support phagocytosis, possibly by acting as an ER-PM MCS tether^24, 25^. More recently, studies have shown that the oxysterol-binding protein-related protein 8 (ORP8) interacts with Sec22b and impacts phagosome maturation^26^. Additional roles for ER-PM MCS in supporting phagocytosis include the phosphatidylinositol transfer proteins Nir2 and Nir3, which function at ER-PM MCS and are required to sustain PI4,5P_2_ levels in the PM to support phagocytosis^27^. Finally, the store-operated calcium channels ORAIs and the ER-resident calcium sensors STIMs are engaged at phagocytosis sites and contribute to stabilizing the ER-PM contact site in addition to increasing cytosolic calcium^28, 29^. Given the complexity of phagocytosis, it is not surprising that MCS play an active role in the engulfment and maturation process.

Phosphotyrosine-based signal transduction is the driver of phagocytosis, which is known to be negatively regulated at least in part by cytosolic SH2 domain-containing phosphatases SHP1 (*PTPN6*) and SHP2 (*PTPN11*)^30^, which bind the inhibitory FcγRIIb receptor through direct binding to its inhibitory immunoreceptor tyrosine-based inhibitory motifs (ITIM) in the cytosolic tail^31, 32^. Given the putative roles of ER-PM MCS in phagocytosis, this raises the possibility that another protein tyrosine phosphatase (PTP) is involved. Specifically, PTP1B, an ER-resident phosphatase whose catalytic domain resides in the cytosol, functions at ER-PM and ER-Endosome MCS^33, 34^. PTP1B can dephosphorylate plasmalemmal and endosomal substrates, such as the insulin and epidermal growth factor receptor^35–37^. PTP1B is now recognized as regulating diverse tyrosine kinase pathways, including the colony-stimulating factor receptor, which is necessary for macrophage formation and activation^38^. The spatial organization of PTP1B at MCS enables it to reach across the contact and antagonize PM-localized signaling events while remaining anchored to the ER membrane. Potential substrates and functions of PTP1B at ER-PM MCS during innate immune responses have not been explored. However, the presence of ER-PM MCS at the site of phagocytosis raises the possibility that PTP-1B may attenuate a portion of the phosphotyrosine signaling downstream of FcγR activation.

Here, we demonstrate that the loss of PTP1B has minimal impact on FcγR-mediated phagocytosis but impacts superoxide generation by dephosphorylating Syk. Using an unbiased proteomics approach, we found that the SH2 domain-containing transforming protein 1 (Shc1) is a critical link between Syk and macrophage superoxide production.

## Results

### Actin depolymerization in the phagocytic cup allows for expansion of ER-PM MCS

Actin stress fibers can act as a physical barrier that limits ER-PM MCS formation^39^. However, in general, macrophages lack stress fibers and instead have a highly dynamic meshwork of crossed and branched filaments^40^. Given the robust actin depolymerization during phagocytosis, we wondered if removing this meshwork would allow for new contact sites to form. Using the ER-PM MCS reporter construct term MAPPER and mCh-Lifeact to monitor F-actin, we examined whether this reporter combination would enable us to detect changes in F-actin density and ER-PM MCS in live cells. First, we examined these reporters in Chinese hamster ovary (CHO) cells that contain actin stress fibers. Indeed, visualizing GFP-MAPPER and mCh-Lifeact in these cells using TIRF microscopy revealed that they are mutually excluded (Fig. 1A, B). Furthermore, we found that these two probes also respond to treatments that impact F-actin assembly. Specifically, we incubated cells with the RhoA activator II, a treatment known to increase stress fiber thickness^41^. As predicted, we found that, compared to the solvent control, the RhoA activation increases mCh-Lifeact in the TIRF plane and decreases GFP-MAPPER (Fig. 1C-E).

**Figure 1.**
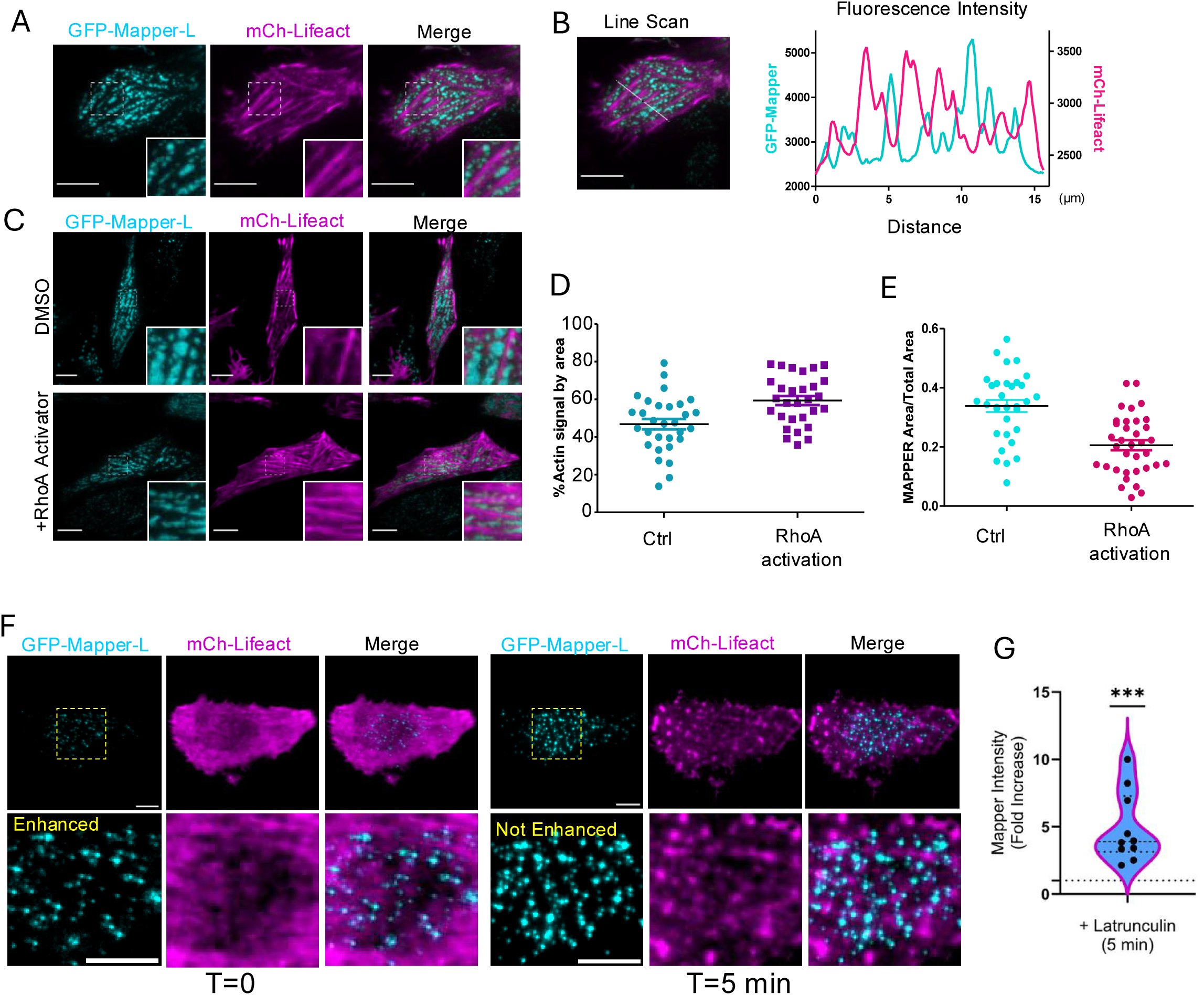
Cortical actin limits ER accessibility to the plasma membrane. **(A)** TIRF microscope images of Chinese hamster ovary cells expressing GFP-MAPPER-L (cyan) and mCherry-Lifeact (magenta). **(B)** Image from panel “A” with a line indicating the position of the line scan and the corresponding histogram. **(C)** Representative TIRF micrograph of CHO cells expressing GFP-MAPPER-L and mCherry-Lifeact in either control or RhoA activator-treated conditions. **(D)** Area occupied in the TIRFM field by Lifeact labeled F-actin, essentially stress fibers, and **(E)** the area occupied by MAPPER-L in control and Rho Activator treated cells. **(F)** TIRF micrographs of a RAW264.7 macrophage expressing GFP-MAPPER-L and mCherry-Lifeact at time 0 and 5 min after actin depolymerization induced by latrunculin B. Bottom row is the magnified inset. For GFP-MAPPER-L at time 0, the image is contrast-enhanced; at the 5 min timepoint, no such enhancement is required. **(G)** Quantitation of 10 individual movies collected on two separate experiments. For statistical analysis, starting intensities of MAPPER-L in individual movies were set equal to 1, and the data were analyzed using a one-sample t-test and a Wilcoxon test. Default GraphPad indicators for significance levels are used throughout the study; *** *P* < 0.005, ** *P* < 0.01, * *P* < 0.05. Scale Bar = 5 μm.

Next, we examined whether acute actin depolymerization induced by latrunculin B impacts the cortical actin and the recruitment of GFP-MAPPER in RAW264.7 macrophages (or simply, RAW cells). For this experiment, RAW cells were co-transfected with GFP-MAPPER and mCh-Lifeact as in Fig. 1A. RAW cells growing on cover glass did not display stress fibers but did have detectable F-actin (Fig. 1F). In these cells, GFP-MAPPER was detectable but not prominent as in the CHO cells (Fig. 1F, contrast-enhanced inset). Following a 5-minute incubation with latrunculin B, a few changes were readily apparent. First, there is a reorganization of F-actin, as monitored by Lifeact. Instead of a meshwork, several small actin puncta appear (Fig. 1F), reminiscent of previously described actin asters^42^. Concomitantly, there was an increase in GFP-MAPPER signal, apparent without the use of contrast enhancement, that correlated with a nearly 5-fold increase in fluorescence intensity in the TIRF field (Fig. 1G). These experiments demonstrate that both stress fibers, as in CHO cells, and the dynamic fenestrated cortical actin present in macrophages limit the formation of ER-PM MCS.

Having established that cortical F-actin removal can promote ER-PM MCS in macrophages, we next examined changes in actin and ER-PM MCS during frustrated phagocytosis. This model (Fig. 2A) is particularly useful since macrophages attempt to internalize the opsonized coverslip, allowing the relative dynamics to be monitored using TIRF microscopy^43, 44^. In this experiment, RAW cells stably expressing mCh-Actin were transiently transfected with the ER-resident YFP-STIM1. At time = 0, the macrophage has entered the TIRF field and contacted the glass. At this instance, the mCherry signal is evident, but little to no YFP signal is detected (Fig. 2B). In conventional particle engulfment or “3D phagocytosis”, as a macrophage engulfs a particle, actin-rich pseudopods wrap around the target while the F-actin at the base of the phagocytic cup is removed^45, 46^. In frustrated phagocytosis, as the cell spreads radially, the ventral surface center is comparable to the base of the phagocytic cup^46^. In the time-lapse, 150 seconds after the initiation of frustrated phagocytosis, actin begins to clear near the initial contact with the opsonized cover glass. At the same time, the progressing membrane remains rich in F-actin. At this point, YFP-STIM1 begins appearing in the actin clearance zone but not where the actin remains abundant (Fig. 2B, arrowheads). At the 360-second time point, once the actin clearance zone is noticeably larger, substantial amounts of the YFP-STIM1 are present in the TIRF plane. Like the stress fibers in CHO cells and the F-actin network in resting macrophages, the F-actin and ER-PM MCS markers show minimal overlap (Fig. 2B, C).

**Figure 2.**
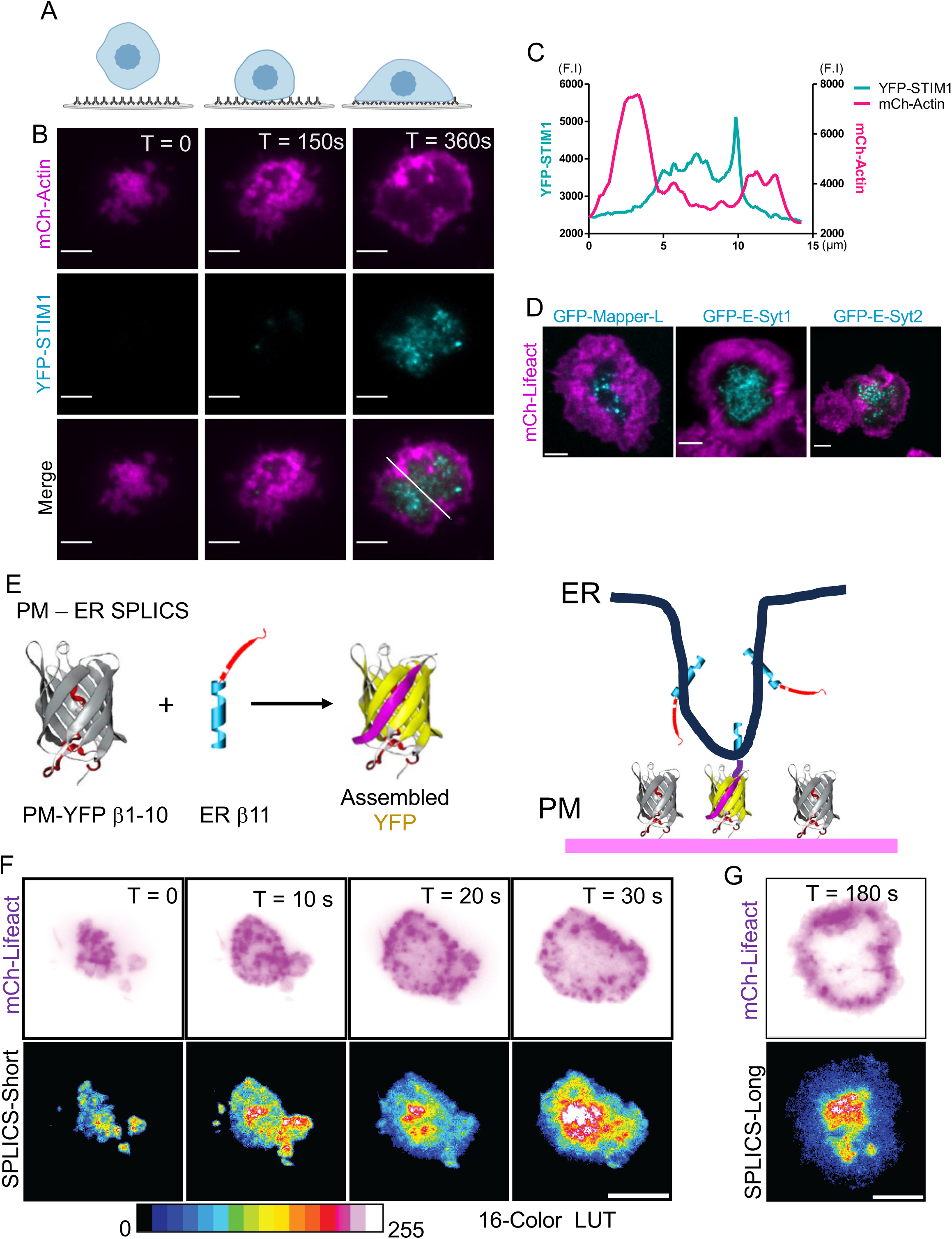
Focal actin disassembly during phagocytosis allows for the formation of ER-PM membrane contact sites. **(A)** Schematic representation of frustrated phagocytosis. Cover glass is opsonized with human IgG and RAW264.7 macrophages are parachuted to initiate frustrated phagocytosis. **(B)** TIRF micrographs at the indicated times of frustrated phagocytosis. RAW264.7 cells stably expressing mCherry-actin (magenta) were transiently transfected with YFP-STIM1 (cyan). The white line in the merged image at 360 sec was used to generate the line scan in **(C)**. F.I. represents fluorescent intensity. **(D)** mCherry-Lifeact expressing RAW 264.7 macrophages were transfected with GFP-MAPPER-L, GFP-E-Syt1 or GFP-E-Syt2 (all pseudo-colored cyan) and were also examined during frustrated phagocytosis. **(E)** Schematic representation of how complementary split GFP/YFP, β1-10 and YFP β11, can be expressed and reformed. Split GFP (or YFP) based contact site sensors (SPLICS) targeting the PM and ER, respectively, to monitor formation of new ER-PM contact sites. RAW 264.7 macrophages were electroporated to express mCherry-Lifeact with either SPLICS-ER-PM Short (**F**) or SPLICS-ER-PM Long (**G**) and monitored using TIRF microscopy during frustrated phagocytosis. The 16-color LUT is used for YFP signal. Scale bar 5 μm.

Since the store-operated calcium channel STIM1-ORAI is known to be activated during phagocytosis, we examined other markers, including GFP-MAPPER and extended synaptotagmin (E-Syt) 1 and 2, two proteins that localize to ER-PM MCS and have both tethering and lipid-transport function^20, 47, 48^. As depicted in Fig. 2D, comparable patterns are seen with each of the GFP-tagged constructs being present in the actin cleared regions but not in the areas equivalent to advancing pseudopods. These results highlight that the coordinated depolymerization and removal of F-actin during phagocytosis provides an opportunity for ER-PM MCS residents to enter the TIRF plane and approach the PM.

MCS are defined as regions of proximity in which the two component organelles come into contact, typically separated by 10-30 nm. The TIRF microscopy has a penetration depth of 100 nm. Thus, ER resident proteins can enter the TIRF plane without entering or forming an MCS. Thus, to complement the TIRF microscopy, we next used a bimolecular fragment complementation assay to demonstrate the formation of these new MCS. Specifically, we chose the SPLICS (SPLit-YFP-based Contact Site sensor) ER-PM reporter developed by Cali and colleagues^49^. In this reporter system, the split YFP β1-10 is targeted to the PM, and the complementary YFP β11 strand is in the ER (Fig. 2E). Two versions are available: a SPLICS-Long that detects contacts of ∼40 nm, and a SPLICS-Short that detects regions where the PM and ER are within ∼10 nm^49^. Once the constructs are in proximity, the YFP molecule can assemble and become fluorescent^50^. We transiently transfected RAW cells with mCh-Lifeact and either SPLICS-long or SPLICS-short and again conducted frustrated phagocytosis experiments. For this, we had to survey the coverslips for cells with minimal YFP signal before engaging the coverslip. As with the other ER-PM MCS resident proteins, we found that most of the SPLICS fluorescence intensity is in the actin-clearance zone, corresponding to the base of the phagocytic cup. Similar results were observed using either SPLICS-Short (Fig. 2F) or SPLICS-Long (Fig. 2,G) supporting the notion that these are *bona fide* ER-PM MCS. Collectively, these results support the idea that the removal of F-actin during phagocytosis progression allows for the establishment of new or the expansion of existing ER-PM MCS.

### PTP1B colocalizes with Fcγ Receptor in the actin clearance zone

PTP1B is an ER-anchored protein tyrosine phosphatase known to function at ER-PM and ER-Endosome MCS. Thus, we wondered if PTP1B was recruited to ER-PM MCS during frustrated phagocytosis. We transiently co-transfected cells with the phagocytic ITAM-containing FcγRIIα-GFP and either mCh-PTP1B or the substrate-trapping mCh-PTP1B^D181A^ and examined the cells after 10 min of frustrated phagocytosis. To visualize F-actin, cells were fixed with paraformaldehyde, permeabilized, and stained with iFluor647-phalloidin. Similar to the other ER-PM MCS resident proteins, we found that both the wild-type and substrate-trapping PTP1B chimeras were present in the actin clearance zone with minimal signal from the actin-rich regions of the membrane (Fig. 3A, B). We occasionally noticed that the pattern of the FcγRIIα-GFP was different in the cells expressing wild-type PTP1B compared to the substrate-trapping mutant. This raised the possibility that PTP1B^D181A^ could engage and trap the phosphorylated FcγRIIα. Indeed, our colocalization analysis of the ventral surface demonstrated that revealed that the substrate-trapping mutant had a higher Pearson’s correlation coefficient than wild-type PTP1B (Fig. 3C). We next sought to determine if the phosphorylation of the FcγRIIα-GFP was required for this colocalization. We again performed frustrated phagocytosis-TIRF microscopy experiments with the substrate-trapping mutant co-expressed with wild-type or ITAM-deleted FcγRIIα-GFP in cells that also express endogenous untagged FcγRs (Fig. 3D). We readily detected overlap between FcγRIIα and PTP1B in the actin clearance zone. However, the overlap was less apparent with the receptor lacking the ITAM. We next wanted to determine whether PTP1B interacted with the phosphorylated FcγRs. To do this, we used the Sleeping Beauty transposon system to generate stable doxycycline-inducible GFP, GFP-PTP1B, and GFP-PTP1B^D181A^ RAW cells to allow for controllable expression^51, 52^. To achieve robust activation of the FcγRs we treated the cells with heat-aggregated IgG (AgIgG), a potent multivalent ligand that can engage and activate multiple FcγRs^53^. After a 10-minute incubation with AgIgG, cells were lysed, and the lysates were incubated with GFP-nanotrap to immunocapture the GFP constructs. As depicted in Figure 3E, the GFP and both GFP-PTP1B isoforms were captured, with only modest amounts of degradation products detected. To our surprise, we did not detect any FcRγ. This finding suggests that, even with a substrate-trapping mutant, the interaction between PTP1B and FcγRs is transient, or that PTP1B interacts with a bridging molecule.

**Figure 3.**
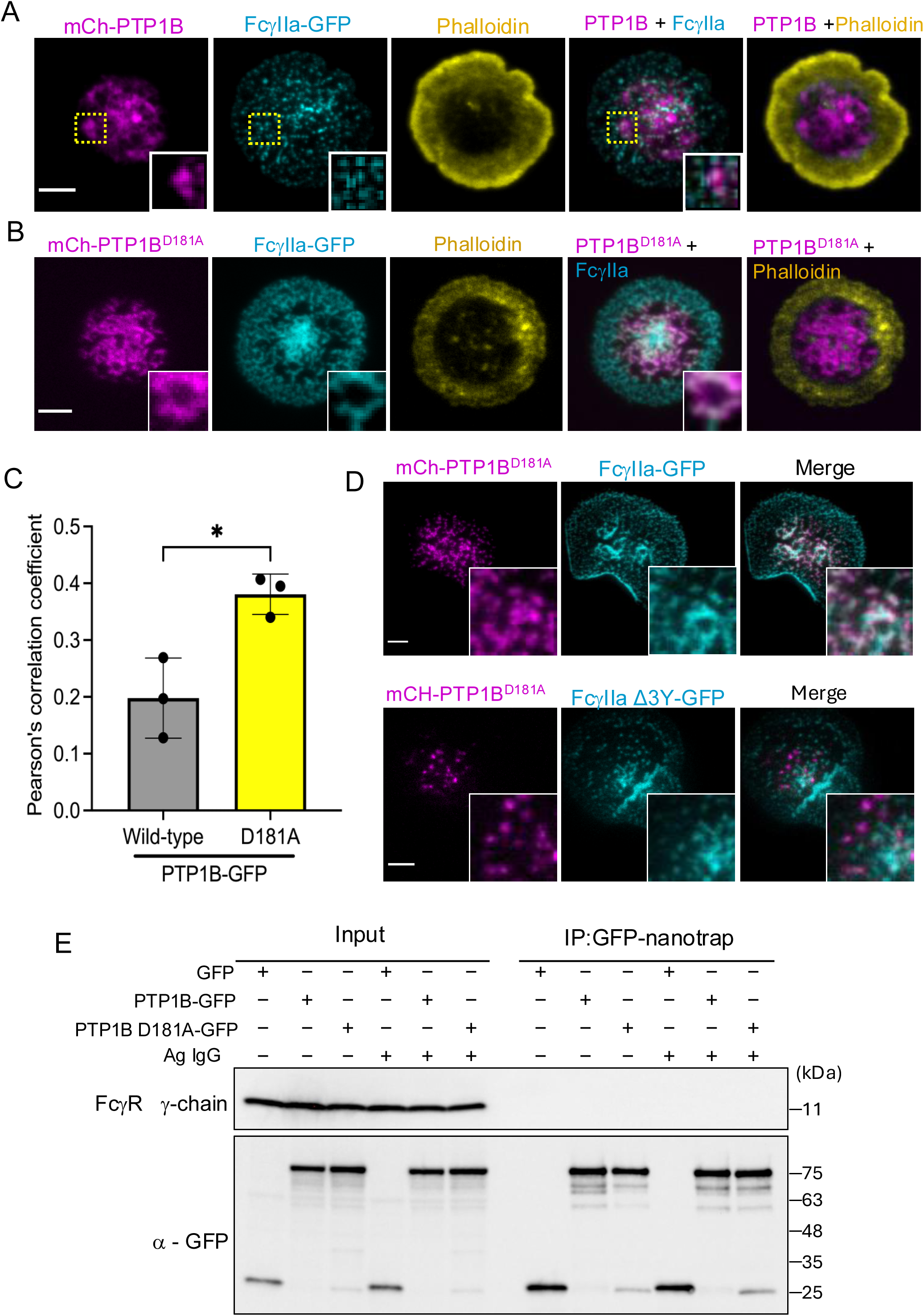
PTP1B co-localizes with FcγR in the actin-cleared zone. **(A)** TIRF micrographs of paraformaldehyde fixed RAW 264.7 macrophages transiently transfected to express PTP1B-mCherry (magenta) and FcγRIIa-GFP (cyan) and stained with iFluor647 conjugated phalloidin (yellow). **(B)** the same as “A” except cells express the substrate trapping mutant mCherry-PTP1B^D181A^. **(C)** Pearson’s colocalization coefficient for FcγRIIa-GFP and either wild-type or substrate trapping D181A PTP1B-mCherry. Mean ± SD and analyzed by an unpaired *t*-test of individual values across n = 3 independent experiments, * = *P* < 0.05. **(D)** TIRF micrographs of paraformaldehyde fixed RAW 264.7 macrophages transiently transfected to express PTP1B^D181A^-mCherry (magenta) and FcγR2a-GFP or FcγR2a lacking amino acids 280-307, which constitute the ITAM, FcγR2a Δ3Y (cyan). **(E)** Representative co-immunoprecipitation immunoblots of stably transfected doxycycline-inducible RAW 264.7 cells transfected with the indicated GFP constructs treated with PBS or aggregated IgG for 10 min. Immunocapture was performed using GFP-Trap and blots probed for the gamma chain of Fc receptor or with an anti-GFP antibody. Scale Bar = 5 μm.

### PTP1B interacts with and attenuates Spleen Tyrosine Kinase

The phosphorylated ITAMs serve as docking sites for Syk kinase^54^, a protein that is also phosphorylated on multiple tyrosine residues^55^, making it a putative PTP1B substrate. We again used frustrated phagocytosis with RAW cells transfected with GFP-Syk and mCh-PTP1B^D181A^ and examined the cells using TIRF microscopy. As depicted in Fig. 4A, Syk and the substrate-trapping mutant were colocalized in the frustrated phagosome. We next used the GFP-nanotrap beads to determine if immunocapture of PTP1B could co-purify Syk. In contrast to the FcγRs, we found that both wild-type and substrate-trapping mutant PTP1B could pull down Syk (Fig. 4B). These results suggest that Syk may be the relevant target of PTP1B in phagocytosis.

**Figure 4.**
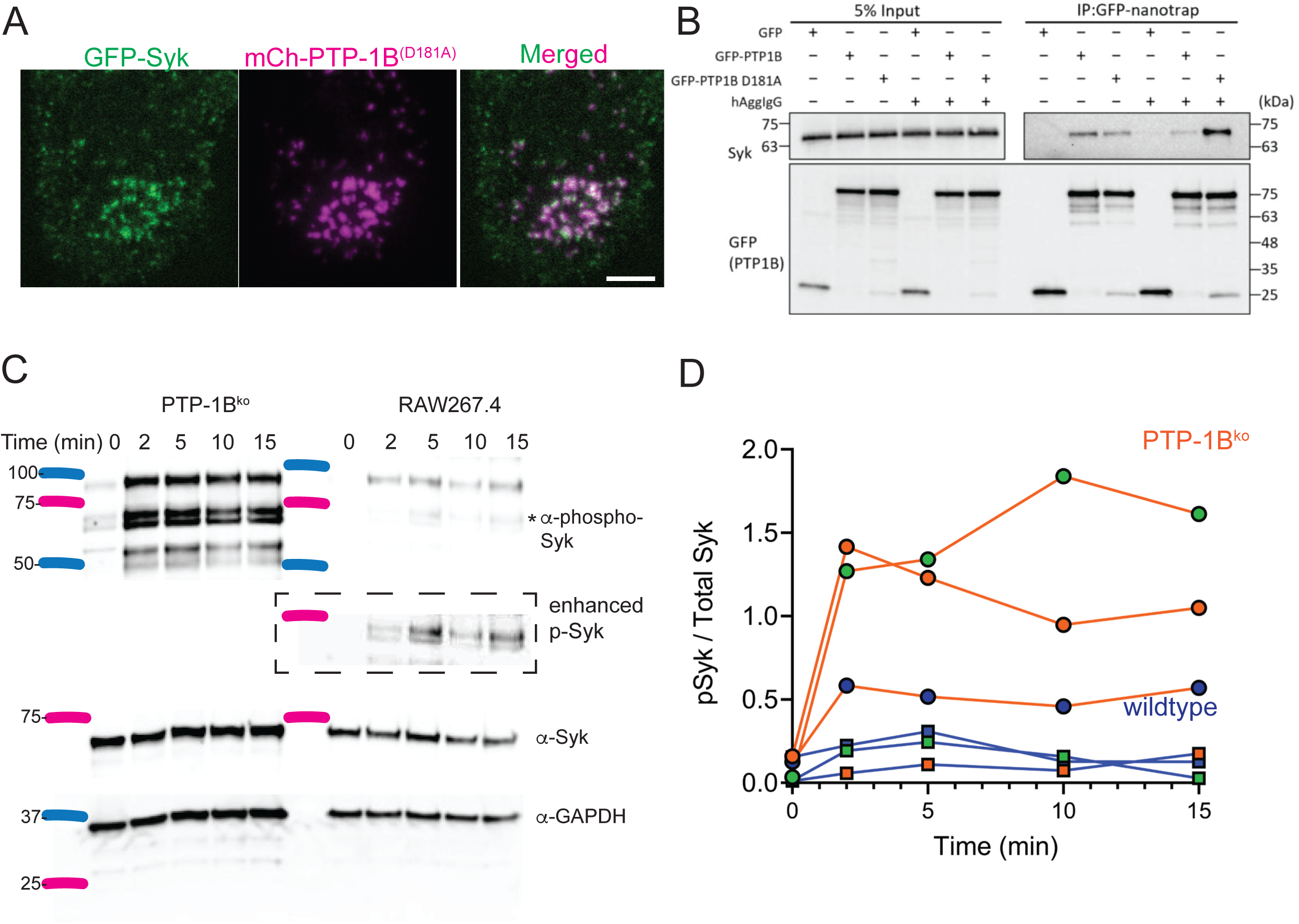
PTP1B interacts with and attenuates phospho-Syk in response to FcγR activation. **(A)** TIRF micrographs of paraformaldehyde fixed RAW 264.7 macrophages during frustrated phagocytosis transiently transfected to express PTP1B^D181A^-mCherry (magenta) and GFP-Syk. Scale Bar = 5 μm. **(B)** Representative co-immunoprecipitation immunoblots of doxycycline-inducible RAW264.7 cells, as in Fig. 3E, treated with PBS or aggregated IgG for 10 min. Immunocapture was performed using GFP-nanotrap, and blots were probed for endogenous Syk and anti-GFP antibody. **(C)** Parental and PTP1B^ko^ RAW 264.7 macrophages were incubated with aggregated IgG for the indicated times. The dotted box highlights a contrast-enhanced version of the panel above since the PTP1B knockouts have such intense bands. **(D)** Line chart of three individual experiments depicted in **“C”.** Given variability in the magnitude of the PTP1B^ko^ cell response, the mean of the orange lines (PTP1B^ko^) is not significantly different from the mean of the blue lines *(P* ∼ 0.10).

To determine if PTP1B was influencing the degree of phosphorylation of Syk, we used CRISPR-Cas9 to generate PTP1B knockout clones. For the next experiment, we conducted a time course using AgIgG as the stimulus and determined phospho-Syk and total Syk levels in cell lysates by immunoblotting. As depicted in Fig. 4C at time = 0, there is minimal p-Syk in the wild-type and PTP1B^ko^ cells. However, the addition of AgIgG led to a modest and sustained increase in p-Syk in parental cells and a substantial increase in PTP1B^ko^ cells (Fig. 4D). These results suggest that PTP1B not only physically interacts with Syk but also dephosphorylates it downstream of FcγR activation.

### Loss of PTP1B and hyperactive Syk increase superoxide production

A variety of CRISPR-based genetic screens using THP1 and U937 cells failed to identify a role for PTP1B in regulating numerous phagocytic pathways^55–57^; nevertheless, we decided to examine its potential role in RAW macrophages. First, we conducted a time-course experiment again monitoring phospho-Syk during the phagocytosis of IgG-opsonized sheep red blood cells. As depicted in Fig. 5A and 5B, the synchronized addition of IgG-SRBCs increased phospho-Syk, plateauing around 2 min and then decreasing over time. Similarly, the PTP1B^ko^ RAW cells show minimal phospho-Syk at rest, reach a plateau at 2 mins, and then decrease. However, in this situation, the total p-Syk levels across all time points are ≍4-fold higher than those in wild-type cells. Thus, as with AgIgG stimulation, Syk is negatively regulated by PTP1B during FcγR-mediated phagocytosis.

**Figure 5.**
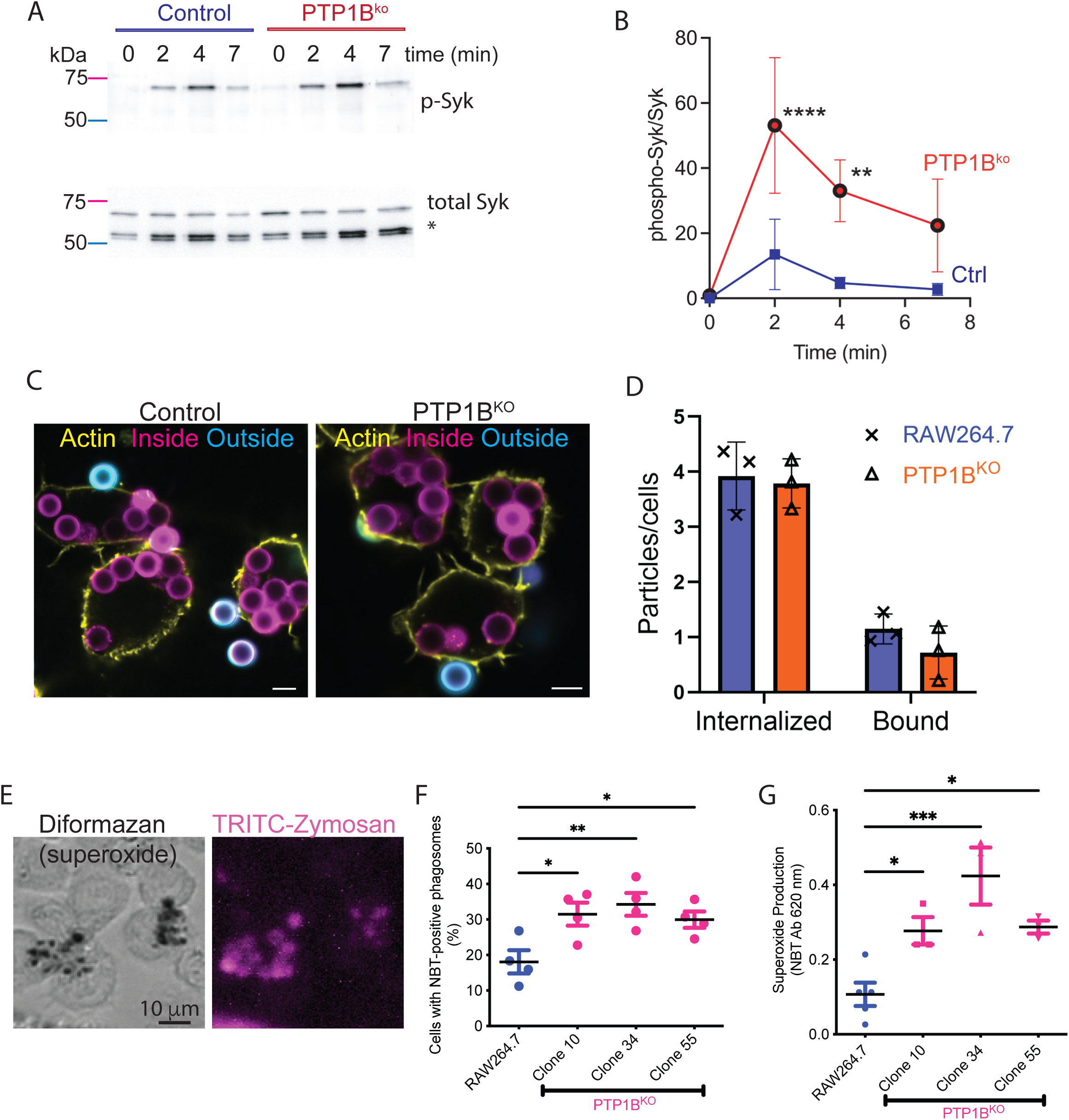
Loss of PTP1B results in increased superoxide production without altering phagocytic efficiency. **(A)** Representative immunoblot of a time course of RAW 264.7 cells incubated with IgG-opsonized sheep red blood cells (sRBC). **(B)** Quantitation of “A”. Data represent the means ± SD from 4 biological replicates. *** *P* < 0.005 and ** *P* < 0.01 by 2-way ANOVA with Bonferroni post-hoc test. **(C)** Confocal micrographs of parental and PTP1B^ko^ RAW 267.4 cells after 10 min incubation with human IgG-opsonized latex beads. Cells are delineated with iFluor 647 phalloidin, all beads are labeled with rhodamine (magenta), and outside beads are labeled with Alexa 488 conjugated Donkey anti-human IgG. Scale Bar = 5 μm. **(D)** Quantitation of “C” from n=3 biological replicates. Data are the mean ± SD. Results are non-significant using an unpaired *t*-test. **(E)** Brightfield and fluorescent micrographs are RAW 264.7 macrophage 10 min post addition of TRITC-conjugated zymosan particles incubated with nitroblue tetrazolium. **(F)** Quantitation of NBT-positive phagosomes in parental and three PTP1B^ko^ clones. Data are the mean ± SD from n=4 biological replicates. * *P* < 0.05, ** *P* < 0.01 as determined by one-way ANOVA and Tukey’s post-hoc test. **(G)** Quantitation of superoxide production in phagosomes following NBT solubilization and measurement of OD 630nm. Data are the mean ± SD from n=4 biological replicates. * *P* < 0.05, *** *P* < 0.005 as determined by one-way ANOVA and Tukey’s post-hoc test.

We next wondered whether the prolonged phospho-Syk levels would affect the extent and efficiency of phagocytosis. Thus, we used IgG-latex beads as prey and monitored phagocytosis in the wild-type and PTP1B^ko^ cells. As shown in Fig. 5C and 5D, we did not detect any appreciable difference with this phagocytic target, consistent with previous CRISPR screens^55–57^. Although we did not observe a change in the degree of particle engulfment, we wondered whether another critical downstream pathway might be affected. Specifically, the ability of macrophages to activate the NADPH oxidase 2 (NOX2) complex and generate antimicrobial superoxide. To examine this possibility, we used nitroblue tetrazolium, a membrane-permeant reagent that specifically reacts with superoxide to form a dark precipitate called diformazan. Since red blood cells are already dark in color and latex beads can refract light, we used IgG-opsonized zymosan, as we have previously^58^. As seen in Figure 5E, many of the TRITC-zymosan-containing phagosomes also contain the diformazan. In these experiments, we compared parental RAW cells with three PTP1B^ko^ clones and counted the number of diformazan-positive phagosomes. We previously reported that not every phagosome in RAW macrophages produces superoxide^58^, and this observation was consistent in the current study. As depicted in Figure 5F, all three of the knockout clones had more difromazan-positive phagosomes than the parental cells, indicating more active NOX2 complexes. We next used a more quantitative measure to assess superoxide production across the entire population^59^. To do this, we again used nitroblue tetrazolium – diformazan conversion, but this time at the end of the experiment, we solubilized the diformazan and measured its absorbance at 620 nm^59^. As illustrated in Fig. 5G, all three knockout clones produced more superoxide than the parental RAW cells. These results demonstrate that enhanced phospho-Syk in cells lacking PTP1B contributes to increased superoxide production.

### DAG-based signaling is not enhanced in PTP1B-deficient RAW macrophages

Syk is known to have many downstream targets but lacks a direct link to the NOX2 complex, which is mainly regulated by phospho-Serine based signals^60–62^. However, Syk is known to phosphorylate phospholipase C isoforms^63, 64^, which, in turn, hydrolyze PI4,5P_2_ to generate diacylglycerol (DAG) in the phagocytic cup. This DAG activates protein kinase C isoforms^65, 66^, which phosphorylate NOX2 components, thereby promoting assembly and activation^60^. Thus, we wanted to know whether enhanced phospho-Syk could lead to increased PKC α/β activation thereby explaining the increase in superoxide production. To examine this, we again conducted a time-course analysis of cells treated with AgIgG and probed with a broad-spectrum anti-PKCα/β substrate antibody. As illustrated in Fig. 6A and B, numerous bands are detected in both the parent and PTP1B^ko^. However, no apparent differences were seen in the two cell types. We previously showed that DAG kinase β rapidly converts DAG to phosphatidic acid, thereby limiting the actions of DAG and PKC in phagocytosis^58^. Thus, it would seem that the DAG-PKC signaling does not explain the enhanced superoxide production in the PTP1B^ko^ cells.

**Figure 6.**
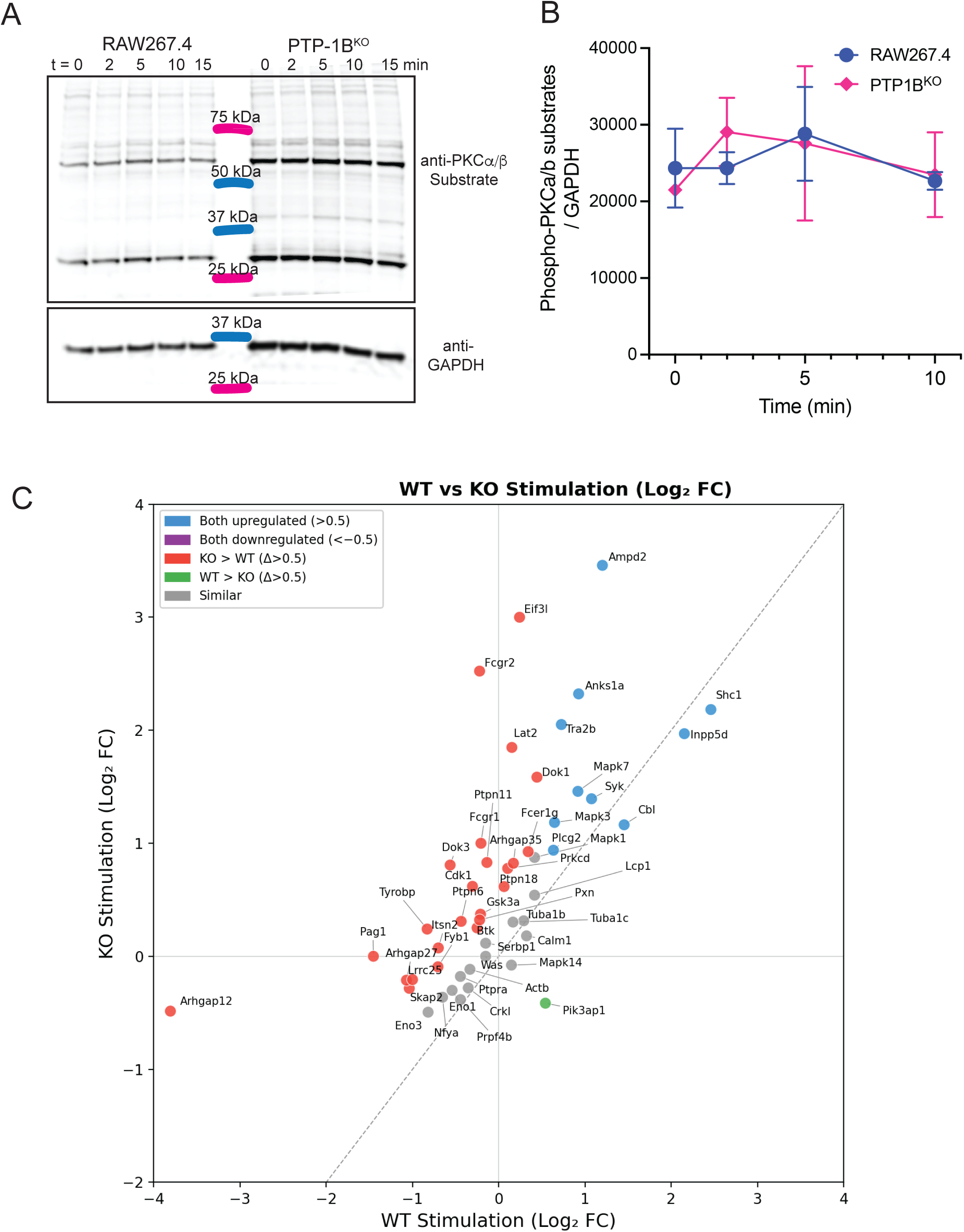
Protein kinase C activation and alterations in phospho-tyrosine signaling following aggregated IgG treatment. **(A)** Representative immunoblots monitoring the increase in the phosphorylation of Protein Kinase C α and β isoforms following the addition of aggregated IgG for the indicated amounts of time. **(B)** Quantitation of “A”. Data are the mean ± SD from n=4 biological replicates. **(C)** Scatter plot comparing phosphotyrosine (pY) signaling responses to stimulation in wild-type (WT) and KO conditions. Each point represents an individual phosphoprotein (top 50 hits depicted). The x- and y-axes show the WT and KO stimulation responses, respectively, expressed as Log fold change (Log FC) relative to unstimulated controls. The dashed diagonal indicates equal fold change between conditions. Points are colored by classification: proteins upregulated upon stimulation in both conditions (Log FC > 0.5 in both; blue), proteins with a greater KO response than WT (KO − WT Log FC > 0.5; red), proteins with a greater WT response than KO (WT − KO Log FC > 0.5; green), and proteins with similar responses between conditions (gray). Full results available in Supplemental Table 1.

### p52Shc is tyrosine phosphorylated downstream of Src family kinase and Syk

We next sought to determine the link between phosphorylated Syk and NOX2 activation. Thus, we used an unbiased proteomics approach to identify proteins with phosphotyrosine (pY) residues that increased during IgG-mediated phagocytosis. To do this, we used a super-binder SH2 domain to enrich for pY peptides coupled with mass spectrometry^67, 68^. This approach identified a variety of proteins relevant to IgG-mediated phagocytosis including Syk, PLCγ2 and the E3-ligase Cbl (Fig. 6C and Supplemental Table 1). Many additional pY proteins were not responsive to AgIgG stimulation. Analysis of the top 50 proteins by aggregate pY signal across two biological replicates highlighted Syk and Shc1 based on their high abundance and robust stimulation-induced responses.

Restricting analysis to the top 50 pY proteins by aggregate signal—thereby focusing on the most robustly detected species—Syk ranks first by total pY abundance, making it the dominant phosphotyrosine protein in the dataset (Fig. 6C). Within this high-signal subset, Syk also exhibits a reproducible stimulation-induced increase in both WT and KO cells (mean log2FC ∼1.3 in each genotype), with consistent directionality across biological replicates. Shc1, which ranks sixth by total pY signal, displays the largest positive WT stimulation response within the set of top 50 pY proteins (mean WT log2FC ≍2.5–2.6, ≍6-fold increase) and remains strongly inducible in the PTP-1B KO (rank 4 for KO stimulation response). Global replicate concordance supports the robustness of these observations: raw spectral counts show moderate-to-strong agreement between biological replicates (Spearman *ρ* ≈ 0.6–0.76 across conditions), and stimulation-induced log2 fold changes show consistent directional reproducibility (Spearman *ρ* ≈ 0.5). While spectral counting reflects peptide detection frequency rather than absolute phosphorylation stoichiometry—and individual phosphopeptides vary in ionization and MS detectability—stimulation-dependent changes within proteins are reproducible across replicates. Together, these findings indicate that Syk stands out by absolute tyrosine phosphorylation and reproducible inducibility, whereas Shc1 stands out by the magnitude of stimulation-dependent phosphorylation within the high-confidence, high-signal subset.

Shc1 (SH2 domain-containing transforming protein 1) is an adaptor protein that plays a vital role in cellular signaling pathways, particularly those involving receptor tyrosine kinases^69^. Shc1 contains several protein-protein interaction domains, including SH2, collagen homology, and PTB (phosphotyrosine binding) domains (Fig. 7A). Shc1 is expressed as three splice variants, p66, p52, and p46, with p66 being the best characterized for its roles in EGF and insulin signaling^70^. Importantly, evidence has recently demonstrated that the p52 isoform can support the activation of NOX2 in aged hepatocytes^71^, whereas silencing p66 Shc1 in RAW cells reduces superoxide production in macrophages by ≍30%^72^. Thus, phospho-Shc1 appears to be the missing link between Syk and NOX2 activation.

**Figure 7.**
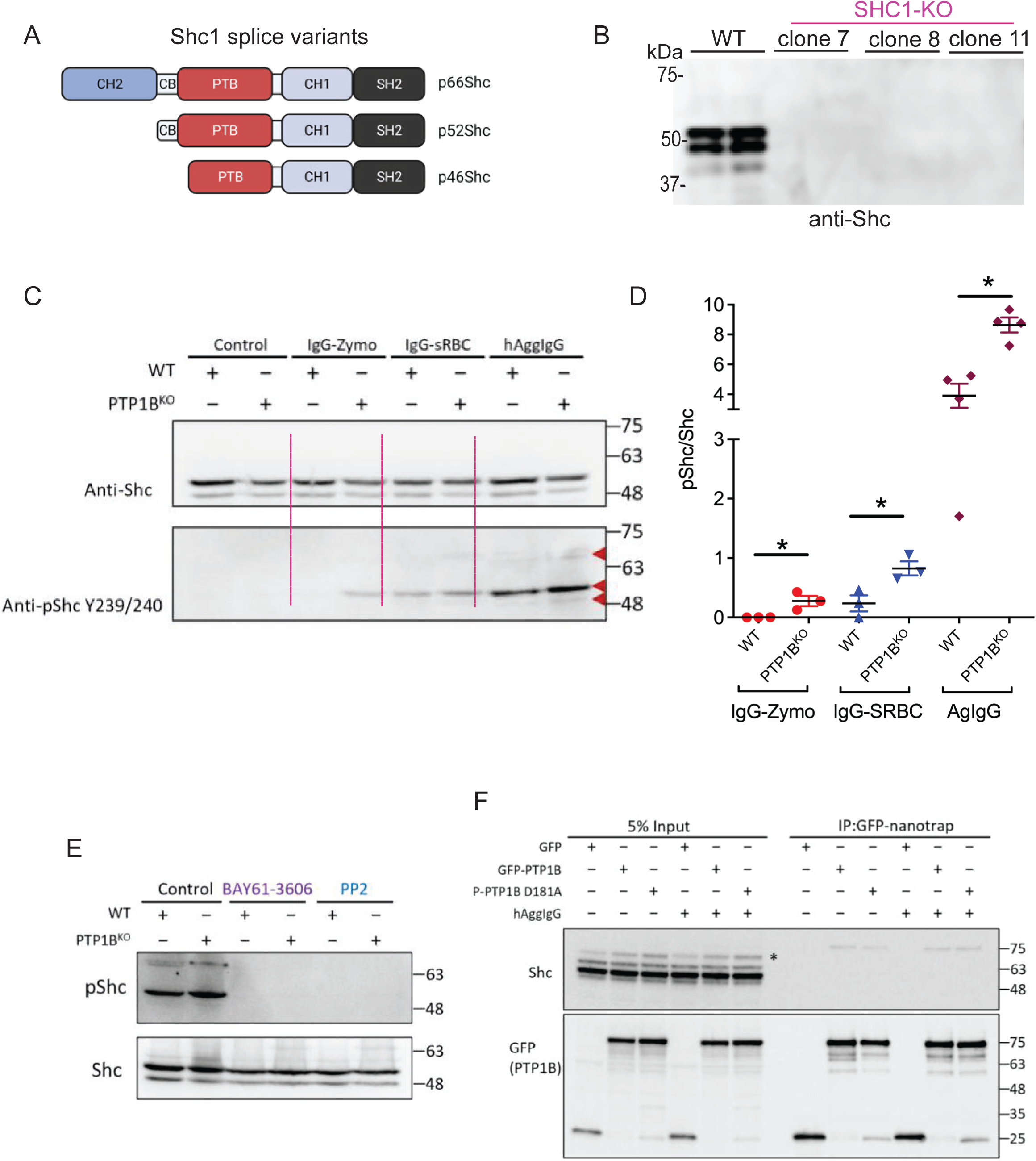
p52SHC1 is phosphorylated during phagocytosis downstream of SFKs and Syk. **(A)** Schematic representation of the three primary isoforms of SHC1, p66, p52 and p46. PTB, phosphotyrosine-binding domain, SH2, Src homology domain 2, CB, cytochrome c binding domain, and CH1 and CH2, collogen homologous region 1 and 2. **(B)** immunoblot of parental RAW264.7 and SHC1^ko^ cells probed with an anti-SHC1 antibody. The bands corresponding to p52-SHC1 and p46-SHC1 are absent from the three knockout clones. **(C)** Immunoblots of parental and PTP1B^ko^ cells treated with IgG-opsonized zymosan, sheep red blood cells, or aggregated IgG probed with either anti-SHC1 or anti-phospho-SHC1. Red arrowheads indicate the expected positions for p66, p52 and p46, respectively. **(D)** Quantitation of “C”. Data represent the mean ± SD. * *P* < 0.05 using an unpaired *t*-test. **(E)** immunoblots using an anti-phospho-SHC1 antibody in parental and knockout cells stimulated for 5 min with aggregated IgG in the presence or absence of Src family kinase inhibitor (PP2) or the Syk inhibitor (BAY61-3606). **(F)** Representative co-immunoprecipitation immunoblots of doxycycline-inducible RAW264.7 cells, as in Fig. 3E, treated with PBS or aggregated IgG for 10 min. Immunocapture was performed using GFP-nanotrap, and blots were probed anti-GFP and anti-SHC1 antibodies. The GFP blots provided for reference are re-used from Fig. 3E.

Using CRISPR-Cas9, we knocked out all three isoforms of Shc1. Immunoblotting confirmed the antibody’s specificity and showed that p52 and p46 are the dominant isoforms, with little p66 expressed in our parental or knockout RAW cells (Fig. 7B). We next examined Shc1 phosphorylation in response to the three FcγR ligands used in the preceding experiments —IgG-Zymo, IgG-SRBCs, and AgIgG — in both parental and PTP1B^ko^ cells. As depicted in Fig. 7C and D, despite varying stimulation magnitudes, the PTP1B^ko^ cells had 2-fold more phospho-52-Shc1 than the parental cells. We also examined the dependency of this phosphorylation on Src-family kinases (SFK) and Syk by using PP2 (a SFK inhibitor) and BAY61-3606 (Syk inhibitor). In both cases, the inhibitors completely abolished phospho-Shc1 (Fig. 7E). Finally, we want to determine whether Shc1 is a direct substrate of PTP1B. We again used the GFP-PTP1B substrate-trapping mutant together with the GFP-nanotrap to immunocapture PTP1B and any bound proteins. This analysis failed to detect any of the Shc1 isoforms (Fig. 7F). Together, these results are consistent with the notion that Shc1 is phosphorylated downstream of SFKs and Syk and that loss of PTP1B results in increased phospho-Shc1.

### Loss of Shc1 reduces superoxide production

We next examined whether the Shc1^ko^ clones had reduced superoxide production in response to IgG-zymosan. We again used the NBT-diformazan assay as in Fig. 5G and solubilized the diformazan to quantify the superoxide production spectrophotometrically. The three clones showed variable reductions in superoxide production, ranging from ≍25% to ≍60% (Fig. 8A). The previous hepatocyte paper demonstrated that p47phox, a soluble NOX2 component, and Shc1 could physically interact^71^. We were unable to detect interactions by coIP. Instead, we used a proximity ligation assay (PLA) as an alternate readout (Fig. 8B)^73^. In this assay, we quantified the number of PLA puncta in resting cells and in cells undergoing frustrated phagocytosis (Fig. 8C-E). We did not detect any significant differences between the parental and PTP1B^ko^ cells grown under resting conditions on unopsonized cover glass. However, during frustrated phagocytosis, we quantified the PLA puncta throughout the cells (Fig. 8D) or near the cover glass on the ventral side of the cell (Fig. 8E) and found a ≍2-fold increase. These results are consistent with previous studies showing that Shc1 influences NOX2-mediated superoxide production through interactions with p47phox.

**Figure 8.**
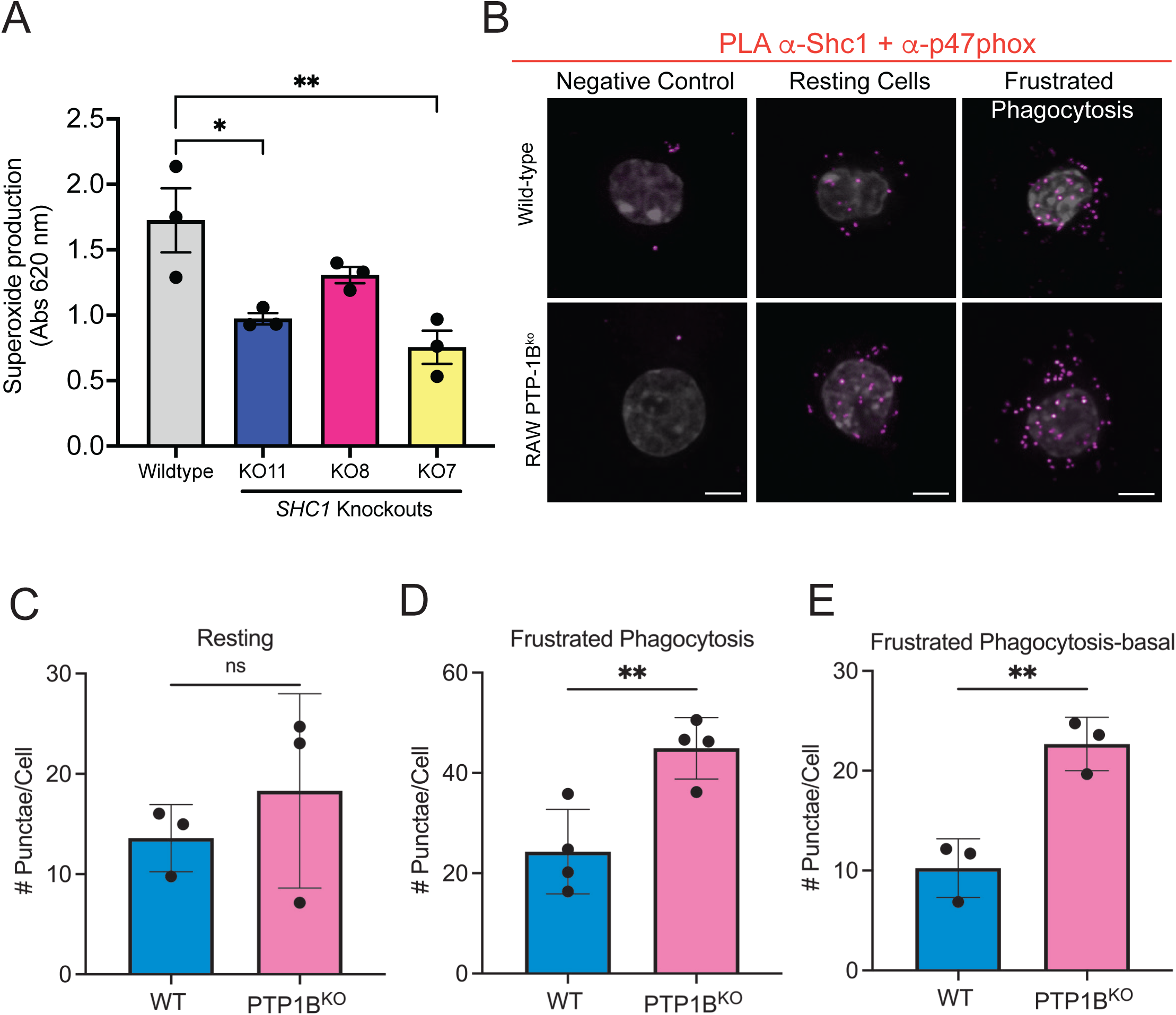
Loss of SHC1 reduces superoxide production. **(A)** Quantitation of superoxide production in parent and SHC1 knockout RAW264.7 cell lines. Macrophages were incubated with IgG-opsonized zymosan for 15 min in the presence of NBT. NBT was solubilized using methanol, and the absorbance was measured at 620 nm. Data represent the mean ± SD from independent biological replicates, n=3. * *P* < 0.05, ** *P* < 0.01 as determined by one-way ANOVA and Tukey’s post hoc test. **(B)** Collapsed Z-slices of parental and PTP1B^ko^ RAW267.4 macrophages grown on cover glass or parachuted onto IgG-coated cover glass and examined using proximity ligation assay. Specifically using a rabbit anti-p47phox and a mouse anti-SHC1. Puncta indicate areas where the antibodies are proximal (magenta), and the nucleus stained with DAPI (70% transparent, white) is included. Scale bar = 5 μm. **(C-D)** Quantitation of the relevant head-to-head comparisons in “B”. Data are the means ± SD from n=3 (C) and n = 4 (D) biological replicates. ** *P* < 0.01 using an unpaired *t*-test. **(E)** Quantitation of the ∼250 nm adjacent to the coverslip. Data are the means ± SD from n=3 (C) biological replicates. ** *P* < 0.01 using an unpaired *t*-test.

## Discussion

Our findings reveal that acute actin depolymerization during phagocytosis creates permissive conditions for the formation of new ER-PM MCS or the expansion of pre-existing ones. The increase in ER-PM MCS enables the recruitment of numerous resident ER-PM MCS proteins, including, but not limited to, ORAI-STIM, E-Syts, and PTP1B (Fig. 2). This provides new insights into the dynamic membrane remodeling and signaling that accompany FcγR-mediated particle uptake.

These observations have important mechanistic implications for understanding the coordination between FcγR signaling and NOX2 regulation during phagocytosis (Fig. 9). Our results demonstrate that phospho-Syk is a substrate of PTP1B and that these proteins interact at the base of the phagocytic cup. Loss of PTP1B results in heightened and prolonged Syk phosphorylation, which ultimately leads to increased superoxide production. Teleologically, it’s unclear why PTP1B would dephosphorylate Syk during phagocytosis. We initially hypothesized that excessive and prolonged Syk phosphorylation may be detrimental for the completion of phagocytosis or subsequent rounds of particle engulfment. However, this did not seem to be the case at least for FcγR-mediated phagocytosis in RAW cells. A previously published CRISPR-based screen using human U937 macrophage and a range of particles, zymosan, myelin, complement-coated RBCs, and IgG-opsonized, found a modest yet significant increase in the uptake of myelin-coated beads, but not for the other phagocytic prey^56^. Furthermore, a recent pre-print also identified PTP1B as a regulator of amyloid beta phagocytosis mediated by Trem2 in microglial cells, with PTP1B deficiency or inhibition leading to a ∼25% increase in microglial phagocytosis through increased Syk signaling^74^. Alternatively, a study using bone marrow-derived mice with a LysM-Cre system demonstrated that loss of PTP1B reduces *Candida albicans* phagocytosis and reactive oxygen production^75^. However, since PTP1B is a negative regulator of colony-stimulating factor signaling and other pathways^38^, these cells may exhibit an altered phenotype that affects phagocytosis, unlike in RAW and U937 cells. Together, the evidence suggests that PTP1B may be a minor contributor to the regulation of phagocytosis, acting in a receptor- and cell-type-specific manner.

**Figure 9.**
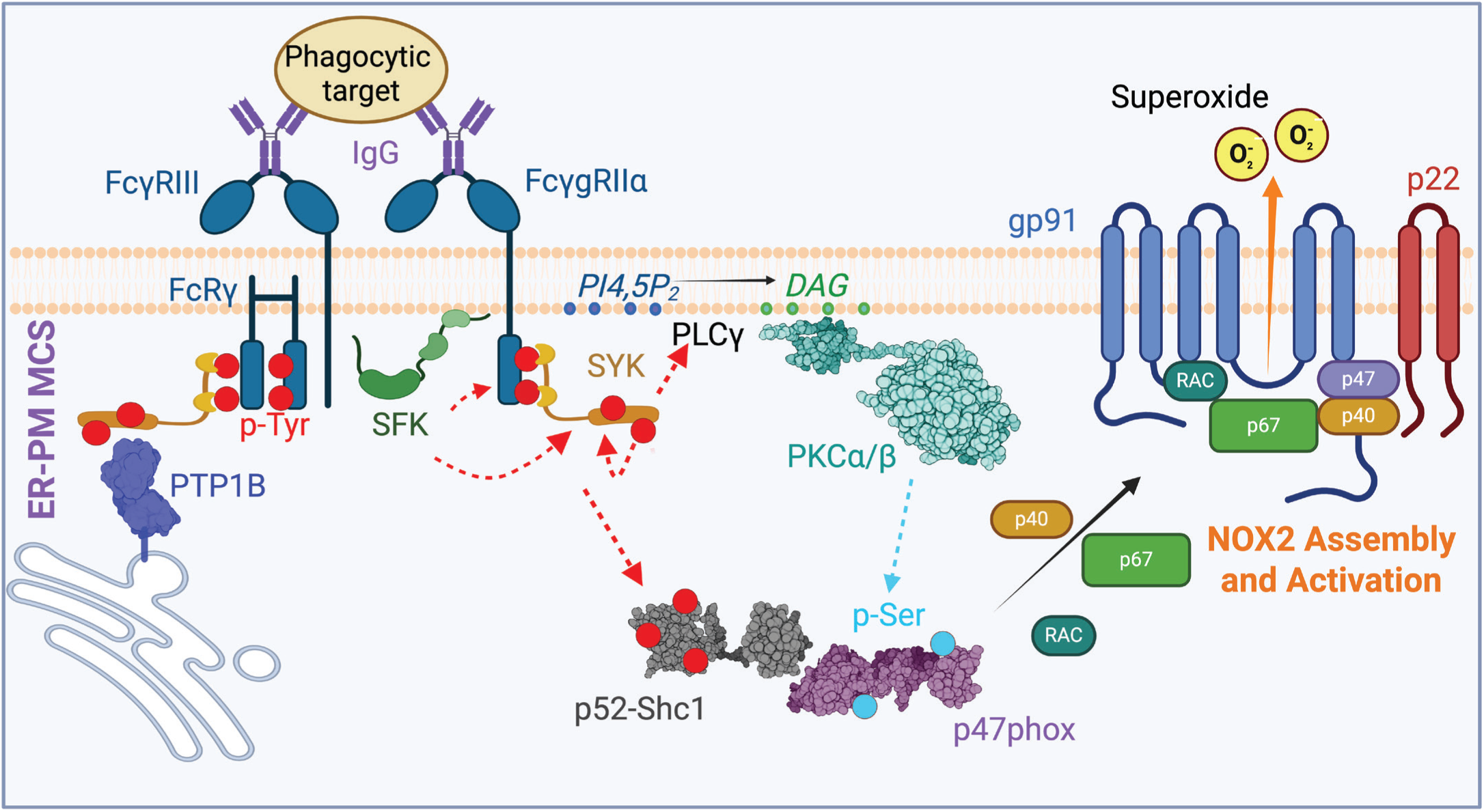
Fcγ Receptor signaling cascade leading to NOX2 activation and negative regulation by PTP1B. Engagement of Fcγ Receptors (FcγRIII and FcγRIIα) by IgG-opsonized phagocytic targets initiates a signaling cascade that culminates in NOX2 assembly and superoxide production. Receptor clustering leads to phosphorylation of immunoreceptor tyrosine-based activation motifs (ITAMs) by Src family kinases (SFKs) (red dotted arrows, tyrosine phosphorylation). Phosphorylated ITAMs recruit and activate Syk kinase, which phosphorylates downstream effectors, including PLCγ (leading to PI4,5P hydrolysis and DAG production) and PKC α/β activation and phosphorylation of NOX2 components (cyan dotted arrow, serine phosphorylation) and the adaptor protein Shc1. Tyrosine-phosphorylated Shc1 interacts with p47phox, facilitating assembly of the NOX2 complex at the membrane. The NOX2 complex comprises the membrane-bound subunits gp91 and p22, along with the cytosolic regulatory subunits p47phox, p67phox, p40phox, and the small GTPase Rac. Upon assembly and activation, NOX2 generates superoxide. The signaling cascade is negatively regulated by protein tyrosine phosphatase 1B (PTP1B), which dephosphorylates Syk at MCS, thereby attenuating the pathway and limiting NOX2 activation. Created in BioRender. Fairn, G. (2026) https://BioRender.com/mc7zbus.

NOX2 activation requires the assembly of multiple cytosolic components (p47phox, p67phox, p40phox, and Rac) with the membrane-bound flavocytochrome b558 complex (gp91 and p22phox) at the plasma membrane/forming phagosome, where it catalyzes superoxide production, which is essential for pathogen killing^76–78^. Given the toxic nature of superoxide, in resting cells, p47phox resides in the cytosol in an autoinhibitory state to limit spurious activation of NOX2. In the resting state, p47phox is maintained in an autoinhibited conformation through intramolecular interactions^79, 80^. The C-terminal autoinhibitory region (AIR) and proline-rich regions physically occupy and block the binding groove formed by the tandem SH3 domains (SH3A and SH3B) within the middle of the protein^81, 82^. This autoinhibition prevents the SH3 domains from binding to their target (the proline-rich region of p22phox). This autoinhibited state also masks the phox homology (PX) domain, preventing it from interacting with phosphatidylinositol 3,4-bisphosphate (PI3,4P_2_) in the membrane^83, 84^. Activation of protein kinase C isoforms is a key activator of p47phox, which has been reported to have eleven Ser residues that can be phosphorylated, with Ser328 and Ser379 being the most critical^61, 85^. This phospho-switch results in the release of the AIR, allowing the PX domain to engage PI3,4P_2_ and SH3 domains to bind the p22phox^79, 80^. Given this sophisticated level of regulation, it is unclear how Shc1supports maximal NOX2 activation. A previous study in aged and palmitate-treated hepatocytes revealed that NOX2 could be activated in these non-phagocytic cells^71^. This study demonstrated that p47phox and Shc1 could physically associate and that phosphorylation of p47phox was reduced in cells lacking Shc1. Although we were unable to co-precipitate Shc1 and p47phox in the RAW macrophages, it is conceivable that transient interactions between the two proteins may allow for relaxation from an auto-inhibited state.

The adaptor protein Shc1 is well established to be recruited to phosphorylated signaling complexes and contribute to Ras/MAPK pathway activation. Our findings, together with previous studies, show that in phagocytic macrophages and non-phagocytic hepatocytes, both p66 and p52 isoforms of Shc1 regulate NOX2 activity. The Syk-Shc1-NOX2 signaling axis we describe here is a critical contributor to superoxide production during phagocytosis. How precisely Shc1 isoforms interact with and regulate p47phox will likely require carefully designed in vitro experiments to complement the growing cell-based literature.

## Materials and Methods

### Reagents and fine chemicals

iFluor 555- and iFluor 647-Phalloidin were purchased from AAT Bioquest (Pleasanton, CA). Latrunculin B was obtained from Abcam (Waltham, MA). 4.2-micron polystyrene beads were purchased from Bangs Labs (Fishers, IN). Rho activator compound CN03 was purchased from Cytoskeleton Inc (Denver CO). BAY61-3606 and PP2 were purchased from Cayman Chemical (Ann Arbor, MI). Nitroblue tetrazolium, human IgG, X-tremeGENE HP were purchased from Sigma-Aldrich (Oakville, Canada). Donkey anti-human secondary antibodies were from Jackson Immunoresearch (West Grove, PA). JetOPTIMUS was obtained from Avantor (Mississauga, Canada). 16% paraformaldehyde and 18 mm round cover glass were purchased from Electron Microscopy Services (Hatfield, PA).

### Cell culture

RAW264.7 and CHO cells were obtained from American Type Culture Collection (ATCC, no.TIB-71TM, CCL-61 TM). RAW 264.7 stably expressing mCherry-Actin was obtained from Dr. Sergio Grinstein (Toronto, Canada) as a kind gift. RAW264.7 cells were cultured in Roswell Park Memorial Institute Medium (RPMI-1640) from Wisent Inc (Saint-Jean-Baptise, Canada) supplemented with heat-inactivated 10% fetal bovine serum (FBS) (Wisent) at 37°C under 5% CO_2_. CHO cells were cultured in DMEM/F12 (Dulbecco’s Modified Eagle Medium/Nutrient Mixture F-12) (Wisent) containing 5% FBS with 5% CO_2_. When cells reached confluency, RAW264.7 cells were washed with warm calcium and magnesium-deficient saline (D-PBS) (Wisent) twice and lifted by gentle scraping. Cells were passaged at 1:10 or 1:5. CHO cells were lifted by Trypsin/EDTA (Wisent) and passaged at 1:20 or 1:10.

### DNA Transfection

RAW264.7 cells were transfected using jetOPTIMUS transfection reagent and X-tremeGENE HP according to the manufacturer’s instructions. Briefly, for JetOptimus reagent, 1 μg of plasmid DNA and 1 μL of JetOptimus were diluted in 100 μL of JetOptimus buffer. The mixture was incubated at room temperature for 10 min, then added to one well of a 12-well plate. For X-tremeGENE HP, 1 μg plasmid DNA was diluted in 100 μL serum-free media, and 3 μL X-tremeGENE HP transfection reagent was added to the following mix. The mixture was incubated at room temperature for 10-15 min, then added to one well of a 12-well plate. Cells were incubated for 18-24 hours post-transfection for further experiments.

### Generation of knockout cell lines

gRNA sequence for murine *PTP1B* (5’-GCCCTTTACCAAACACATGT-3’), *SHC1 (5’-*TGCCAAAGGGGCGACAAGG-3’), and non-targetting (5’-GCACTACCAGAGCTAACTCA-3’) were cloned into pSpCas9(BB)-2A-Puro (PX459) V2.0 following BbsI digestion and T4 DNA ligase mediated ligation as previously described^86^. To deliver the CRISPR plasmid into RAW264.7 macrophages, cells were electroporated using the Neon Transfection System (Life Technologies) as previously described^87^. Briefly, RAW264.7 cells were washed twice with PBS and lifted from the culture flask by gentle scraping. Cells were resuspended in 10% RPMI and centrifuged at 300 xg for 5 minutes. The cell pellet was washed in PBS and centrifuged again to sediment cells. DNA plasmid was delivered by electroporating 5×10^5^ cells with 3-10 μg of plasmid using a single 20-millisecond pulse of 1750V. Transfected cells were selected by puromycin (5 μg/ml) treatment for 72-96 hrs. Dead cells were removed by washing twice with D-PBS, followed by scraping to lift viable cells. Single cells were added to wells of a 96-well plate and allowed to expand in a 50:50 mix of fresh and conditioned RPMI + 10% FBS. Knockouts were identified by western blotting with anti-PTP1B or anti-SHC1 antibodies, respectively.

### Immunoprecipitation

For immunoprecipitation of Syk, RAW264.7 cells were grown in a 6-well plate to 70-80% confluence. Cells were treated with 4 μl of heat-aggregated human IgG^53^ or 100μl of a 10% IgG-opsonized sRBC suspension for the indicated time. The plates were then placed on ice and washed twice with ice-cold PBS with divalent cations. Cells were lysed in ice-cold non-denaturing lysis buffer (20 mM Tris HCl pH 8, 137 mM NaCl, 1% NP-40, 2 mM EDTA) supplemented with phosphatase inhibitor (Roche) and cOmplete EDTA-free Protease Inhibitor Cocktail (Roche). Lysate was transferred to a microcentrifuge tube and incubated at 4°C with constant agitation for 30 min. Cell debris was removed by centrifugation at 12,000 x g for 20 min, and supernatant was collected. GFP-tagged proteins were immunoprecipitated using GFP-Trap magnetic agarose beads (Chromotek) according to the manufacturer’s protocol. Briefly, cell lysates were prepared in lysis buffer and incubated with pre-equilibrated GFP-Trap beads for 1 hour at 4°C with gentle rotation. Beads were washed three times with wash buffer, and bound proteins were eluted by boiling in SDS sample buffer for 10 minutes at 95°C. Samples were then analyzed by SDS-PAGE and western blotting then analyzed samples.

### Western blotting

Protein samples were prepared in SDS Laemmli Buffer and heated at 95 °C for 5 min prior to loading onto a SDS polyacrylamide gel. After running the gel at 150V, the protein was transferred to a PVDF membrane using the Trans-blot Turbo Transfer System according to the manufacturer’s instructions. Membrane was blocked in 5% BSA or 5% Skim milk in TBS-Tween 0.05% for 1 hr, followed by incubation with primary antibody (Anti-phospho-Syk in 1:500; Anti-4G10, Syk, p47phox, gp91, phospho-Ser, SHC, phospho-SHC1 in 1:1000; anti-GAPDH in 1:5000 in 5% BSA TBS-T) overnight at 4°C. Membrane was washed with TBS-T three times for 10 min and incubated with secondary antibody (anti-rabbit IgG-HRP–linked antibody (7074, Cell Signaling Technology) and anti-mouse IgG-HRP–linked antibody (7076, Cell Signaling Technology) at a dilution of 1:5000 in 5% BSA in TBS-T). Membranes were washed three times followed by HRP signal detection using Pierce ECL solution as a substrate (ThermoFisher). Images were captured using the Bio-Rad ChemiDocTM Touch Imaging System or the Azure 600 Imager (Azure Biosystems). As needed, membranes were stripped with Restore Western Blot Stripping Buffer (Thermo Fisher) and reprobed with another antibody.

### Superbinder SH2 pTyr enrichment and mass spectrometry

Affinity purification (AP) of pY-containing peptides was performed essentially as described in Tong et al.^68^ but using a glutathione S-transferase (GST) fusion protein containing two, tandem Src superbinder SH2 domains containing T183V, C188A, and K206L substitutions separated by a 7-residue spacer (EFPGRLE; GST-Src-sSH2.2). GST-Src-sSH2.2 was immobilized on Glutathione MagBeads (GeneScript Catalog No. L00327) under native conditions according to manufacturer’s instructions at a concentration of approx. 2 µg/µl. Next, 20 µl GST-Src-sSH2.2 beads were incubated with approx. 424 nmol tryptic peptides in 0.5 ml AP buffer (50 mM MOPS pH 7.2, 10 mM Na_2_HPO_4_, 50 mM NaCl) at 4 °C for 3 h with gentle inversion mixing. Beads were collected using a magnetic rack and centrifugation at 400 RCF for 1 min, and then washed 3 times with 500 µl of AP buffer. Bound peptides were eluted by three extractions (50 µl each) with 50 mM sodium phenylphosphate in AP buffer. Eluates were pooled (150 µl) and further processed by reduction and alkylation as previously described^68^ followed by digestion overnight at 37°C with trypsin (2 μg; Pierce). Peptides were dried by vacuum centrifugation, desalted on C18 ziptips (Millipore) using a DigestPro MSi (Intavis Bioanalytical Instruments), and dried again by vacuum centrifugation before resuspension in Buffer A (0.1% formic acid). Samples were analyzed by liquid chromatography tandem mass spectrometry using an EASY-nanoLC 1200 system with a 1 h analysis and an Orbitrap Fusion Lumos Tribrid Mass Spectrometer (Thermo Fisher Scientific). The LC portion of the analysis consisted of a 18 min linear gradient running 3-20% of Buffer A to Buffer B (0.1% FA, 80% acetonitrile), followed by a 31 min linear gradient running 20-35% of Buffer A to Buffer B, a 2 min ramp to 100% Buffer B and 9 min hold at 100% Buffer B, all at a flow rate of 250 nL/min. Samples were loaded into a 75 μm x 2 cm Acclaim PepMap 100 Pre-column followed by a 75 µm x 50 cm PepMax RSLC EASY-Spray analytical column filled with 2 µM C_18_ beads (Thermo Fisher Scientific). MS1 acquisition resolution was set to 120000 with an automatic gain control (AGC) target value of 4 x 10^5^ and maximum ion injection time of 50 ms for a scan range of *m/z* 375-1500. Monoisotopic precursor selection (MIPS) was determined at the peptide level with a global intensity threshold of 10000 and only peptides with charge states of 2 to 7 were accepted, with dynamic exclusion set to 10 s. Isolation for MS2 scans was performed in the quadrupole with an isolation window of *m/z* 0.7. MS2 scans were performed in the ion trap with maximum ion injection time of 10 ms, AGC target value of 1 x 10^4^, and higher-energy collisional dissociation (HCD) activation with a normalized collision energy of 30.

MS raw files were analyzed using PEAKS Studio software (Bioinformatics Solutions Inc.) and Proteome Discoverer (version 2.5.0.400), and fragment lists searched against the mouse UniProt Reference database (Uniprot_UP000000589 downloaded Sept. 15, 2020; 21966 entries). For both search algorithms, the parent and fragment mass tolerances were set to 10 ppm and 0.6 Da, respectively, and only complete tryptic peptides with a maximum of three missed cleavages were accepted. Carbamidomethylation of cysteine was specified as a fixed modification; deamidation of asparagine and glutamine, oxidation of methionine, acetylation of the protein N-terminus, and phosphorylation of serine, threonine, and tyrosine residues were specified as variable modifications. Scaffold (version Scaffold_5.3.4, Proteome Software Inc., Portland, OR) was used to validate MS/MS-based peptide and protein identifications. Peptide identifications were accepted if they could be established at greater than 95.0% probability by the Percolator posterior error probability calculation^88^. Protein identifications were accepted if they could be established at greater than 99.0% probability and contained at least 2 identified peptides. Protein probabilities were assigned by the Protein Prophet algorithm^89^. Proteins that contained similar peptides and could not be differentiated based on MS/MS analysis alone were grouped to satisfy the principles of parsimony.

### Confocal microscopy and (Total Internal Reflection Fluorescence) TIRF microscopy

Images were acquired on a Quorum spinning disk microscope system consisting of a Yokogawa X1 head with Borealis, a 63x oil immersion objective with NA of 1.4, and Hamamatsu ImagEM X2 EM-CCD camera. The microscope is equipped with diode-pumped solid-state lasers (405, 488, 552, and 730 nm). Images were also acquired on a Quorum spinning disk with TIRF system consisting of Diskovery/Nipkow spinning disk with dual-pinhole sizes (50 μm & 100 μm), Diskovery TIRF Unit, 63x oil immersion objective with numerical aperture of 1.47, and Hamamatsu ImagEM X2 EM-CCD camera. The system is based on a Leica DMi8 equipped with diode-pumped solid-state lasers (405, 488, 561, and 637 nm). In both microscopes, z-slices encompassing the cell were taken at intervals of 0.2 μm or 0.25 μm. Images were acquired using MetaMorph software and analyzed using ImageJ2/FIJI^90^.

### Immunofluorescence

Cells on a coverslip were fixed in 4% paraformaldehyde for 20 min at room temperature, followed by permeabilization and quenching with 0.1% Triton X-100 and 150 mM glycine for 20 min at room temperature. Coverslips were blocked with 5% BSA in PBS for 60 min, followed by labelling with primary antibody at 1:50 to 1:100 dilutions in 5% BSA for 1 hr. Coverslips were washed twice with PBS for 5 min and incubated with a secondary fluorescent antibody at a 1:1000 dilution in 5% BSA for 40 min. Coverslips were washed three times with PBS for 5 min and imaged with confocal or TIRF microscopy. Where applicable, cells were stained with 1:5000 Phalloidin conjugated with fluorophore for 20 min.

### Nitroblue tetrazolium (NBT) assay

RAW264.7 macrophages were seeded on 18 mm glass coverslip in a 12-well plate and incubated for 18-24 hours. The medium was changed to a fresh complete medium containing 10 μg/ml nitroblue tetrazolium (NBT) for superoxide detection. 40 μl of human IgG-opsonized zymosan was added to each well and incubated for 1hr at 37°C. After the incubation period, cells were washed with PBS and fixed and permeabilized. IgG-Zymosan was probed using tetramethyl rhodamine-conjugated donkey anti-human secondary antibody. Cells with diformazan precipitate (the product of NBT + superoxide) and TMR-zymosan were imaged using the brightfield and TxRED settings on the EVOS Floid imaging system. Cell, phagosome number, and diformazan-positive phagosomes were determined^58^. For quantitative spectrophotometry, 3×10^5^ viable cells were plated and incubated with IgG-zymosan for 1 hour. Cells were washed twice with PBS, once with methanol, and then air-dried. The diformazan deposits in the cells were dissolved by adding 120 μL 2M KOH, followed by 140 μL DMSO with gentle shaking for 10 min at room temperature^59^. The OD_630_ was measured using a BioTek Synergy H1 multimode plate reader.

### Proximity Ligation Assay (PLA)

To monitor the proximity and potential interactions between p47phox and SHC1 in RAW264.7 macrophages, Duolink Proximity Ligation assay kit (Sigma-Aldrich) was used following the manufacturer’s instructions. Briefly, RAW264.7 cells on a coverslip were fixed with 4% paraformaldehyde and permeabilized with 0.1% Triton X-100 in TBS. Cells were blocked with the Duolink blocking solution for 1 hr at 37°C, followed by the incubation of primary antibodies for 1 hr at room temperature. Rabbit polyclonal anti-SHC1 and mouse monoclonal anti-p47phox were used in this assay. A pair of PLA secondary antibody probes conjugated to an oligonucleotide was incubated for 1 hr at 37°C. Ligase was added and incubated for 30 min at 37°C to allow proximity-based joining of the oligonucleotides to form a circular DNA template for rolling-circle amplification. The signal was amplified by incubating the samples with DNA polymerase for 100 min at 37°C, along with fluorescently labeled oligonucleotides that hybridized to complementary sequences within the amplicon. The samples were visualized through confocal microscopy where discrete fluorescent puncta were captured, counted and quantified.

### Image processing, Quantification, and Statistics

Images were acquired using Metamorph and exported as TIFFs. Data analysis and image contrast enhancement were performed using Fiji or ImageJ2 for all experiments. Statistics were performed using GraphPad Prism 10 software. Specific details are included in the figure legends.

## Author contributions

**M.L.** Conceptualization, Methodology, Formal analysis, Investigation, Data Curation, Writing - Original Draft, Visualization. **H.S.Z., M.G., M.L.;** Formal analysis, Investigation. **K.C., L.W.G., M.F.M.;** Methodology, Formal analysis, Investigation. **G.D.F.;** Writing - Review & Editing, Visualization, Supervision, Project administration, Funding acquisition

## Acknowledgments

Figures were generated using Adobe Illustrator 2024-25, FIJI, and BioRender.com. Mass spectrometry was performed at SPARC BioCentre, Hospital for Sick Children (Toronto). The authors would like to thank Kuiru Wei and Janhavi Negwakar for technical assistance in constructing cell lines and with preliminary experiments.

## Additional files

**Supplemental Table 1. Mass Spectrometry results**

